# A novel non-antibiotic selectable marker *GASA6* for plant transformation

**DOI:** 10.1101/2021.07.10.451893

**Authors:** Yuewan Luo, Jiena Gu, Xiaojing Wang, Shengchun Zhang

## Abstract

Selectable markers help the transformed cell/tissue to survive in an otherwise lethal exposure of an antibiotic or herbicide. Unfortunately, almost all the traditional selectable markers are antibiotic and herbicide resistance genes, which are controversial on human health concerns and environmental impact. Novel plant-derived, non-antibiotic, and non-herbicide selectable markers are urgently needed in plant transformation. Our previous work showed that the seedlings of overexpression *Arabidopsis* lines of *AtGASA6* survived on medium with a high concentration of sugar, which leads to the hypothesis that *AtGASA6* could be a selectable marker on media with high or low sugar content. In this study, leaf explants of *AtGASA6* overexpression tobacco lines regenerated shoots on sugar-free shooting medium while those of wild type could not. Moreover, the seeds of *AtGASA6* overexpression tobacco lines germinated and grew into normal seedlings on sugar-free MS medium while those of WT could not. Attractively, no developmental defects were observed in *AtGASA6* transgenic progenies. Using *AtGASA6* as a selectable marker, overexpression tobacco lines of *GAI*, which restrains plant size, were created on sugar-free media. The *GAI* overexpression lines had a smaller plant size than that of control. Considering its plant-derived and non-antibiotic nature, *GASA6* is promising to be used as a selectable marker in plant transformation.

## Introduction

Genetic transformation and genome editing help us create crops with advanced characteristics, such as high yield, high abiotic stress tolerance, and high biotic stress resistance. Efficient selection of transformed cells among a large number of non-transformed cells, which is enabled by selectable markers, is the prerequisite of the production of genetically modified (GM) plants (Yoder and Goldsbrough 1994). The most widely used selectable markers are antibiotic genes that confer resistance to toxins, such as antibiotics and herbicides. These antibiotic genes include *npt II* which encodes the neomycin phosphotransferase II conferring resistance to kanamycin (Fraley et al. 1983), *hpt* that encodes hygromycin phosphotransferase detoxifying hygromycin (Waldron et al. 1985), and *bar* endowing resistance to phosphinothricin (Deblock et al. 1989). Antibiotic genes assist the selection of transformants efficiently. However, whether they harm human health and ecological balance is still unknown. Proteins coded by these antibiotic genes are always undesirable in the final products. In the near future, it is unlikely that transgenic plants using an antibiotic selectable marker will be authorized in the European Union (European directive 2001/18/EC) for both basic research and agricultural application.

Two major strategies were developed in the past few years to reduce the risk of antibiotic selectable markers. One is to remove the selectable marker gene after the initial selection of transformants. This strategy contains numerous approaches, such as co-transformation, site-specific recombination, intrachromosomal homologous recombination, and the transposon-based method (Breyer et al. 2014; Daniell et al. 2001; Ebinuma et al. 1997; Gleave et al. 1999; Goldsbrough et al. 1993; Klaus et al. 2004; Komari et al. 1996; Zubko et al. 2000; Zuo et al. 2001). In these approaches, the target gene will be separated from the selectable marker gene after the initial selection and the selectable marker gene would be removed after one or several generations of crossing. However, these approaches are time-consuming, labor-intensive, and low efficiency. The second strategy is to develop less controversial non-antibiotic selectable markers. Up to date, several non-antibiotic selectable marker candidates have been identified. These candidates include *xylA* (*xylose isomerase*) (Haldrup et al. 1998), *PMI* (*phosphomannose isomerase*), *GALT* (*UDP-glucose:galactose-1-phosphate uridyltransferase*) (Joersbo et al. 2003), *atlD* (*arabitol dehydrogenase*) (LaFayette et al. 2005), *dao1* (*d-amino acid oxidase*) (Erikson et al. 2004; Hattasch et al. 2009), *ilvA* (*threonine deaminase*) (Ebmeier et al. 2004), *dsdA* (*d-serine ammonia lyase*) (Erikson et al. 2005; Lai et al. 2011), *lyr* (lysine racemase) (Chen et al. 2010), *D-LDH* (*D-lactate dehydrogenase*) (Wienstroer et al. 2012) and *MSRB* (*Methionine sulfoxide reductase B*) (Li et al. 2013). Unfortunately, these candidates have some defects. First, most of these candidates are from bacteria, which causes food safety concerns. Second, some of these genes interfere with endogenous metabolic pathways and hence affect the normal growth of transgenic plants. Last, selection reagents for these candidates are too expensive, which hampers the large-scale application. Due to these shortcomings, these candidates have not been broadly used in plant transformation. Up to the present, more than 50% of plant species approved for commercialization contain antibiotic selectable marker genes. It is extremely urgent to develop novel non-antibiotic selectable markers, that are safe, cost-saving, and efficient, in plant transformation.

GASA (gibberellic acid stimulated transcripts in Arabidopsis) family genes are involved in the regulation of seed germination (Rubinovich and Weiss 2010), root formation (Taylor and Scheuring 1994; Zimmermann et al. 2010), establishment of seed size (Roxrud et al. 2007), stem growth and flowering time (Ben-Nissan et al. 2004; Zhang et al. 2009), fruit development and ripening (Moyano-Canete et al. 2013), fiber development (Liu et al. 2013), and leaf expansion (Sun et al. 2013). Our previous work showed that *AtGASA6*, one of the GASA family members from *Arabidopsis*, affected seed germination and responded to gibberellic acid (GA), abscisic acid (ABA), and glucose signaling (Zhong et al. 2015). Overexpression of *AtGASA6* in *Arabidopsis* confers seed germination and seedling growing on the medium with a high concentration of sugar (Zhong et al. 2015). Therefore, we hypothesize that this plant-derived *GASA6* might be used as a non-antibiotic selectable marker in plant transformation on the medium with certain sugar content.

In this work, *AtGASA6* overexpression tobacco was created. The shoot and root regeneration of leaf explants of *AtGASA6* overexpression tobacco on media with high concentration sucrose and without sucrose were investigated. The morphological variations of *AtGASA6* overexpression tobacco were evaluated. Using *AtGASA6* as a selectable marker, *AtGAI* overexpression tobacco lines were generated and the phenotyping of overexpression lines was conducted.

## Material and methods

### Plant material

The *Nicotiana tabacum* variety K326 was used in this work. Seeds were surface-sterilized in 1% sodium hypochlorite solution for 20 min. Then the seeds were rinsed in sterile water 4-5 times and germinated *in vitro* on the MS medium containing 3% sucrose and solidified with 0.8% agar (Murashige and Skoog 1962). Plants grew in a controlled-environmental growth chamber at 24°C and about 70% humidity with a 16-h-light/8-h-dark cycle. Six weeks after germination, 1 cm^2^ leaf discs were isolated from fully expanded leaves and used as explants for transformation.

### Vector construction

To create *AtGASA6* overexpression plants, *AtGASA6* gene (Genbank accession number *At1G74670*) was isolated from *Arabidopsis thaliana* and was subsequently cloned into the vector pBIN19. The PCR primers 5’-CGCTCTAGAATGGCCAAACTCATAACTTC-3’ (forward) and 5’-ATAGGATCCGATGGACATTTTGGTCCACC-3’ (reverse) were used to amplify the coding sequence (CDS) of *AtGASA6*. The CDS of *AtGASA6* was ligated downstream of the *CaMV 35S* promoter in the vector pBIN19 via the XbaI and SacI sites to create the expression cassette CaMV35Spro::*AtGASA6*. In this construct, *nptII* (CaMV35Spro::*nptII*) was used as a selectable marker.

To create *AtGAI* overexpression plants, The CDS of *AtGAI* (Genbank accession number AT1G14920) was inserted downstream of the *CaMV* 35S promoter to generate the overexpression cassette *CaMV*35Spro::*AtGAI*. In this construct, *AtGASA6* (CaMV35Spro::*AtGASA6*) was used as a selectable marker.

### Plant transformation and regeneration to create *AtGASA6* overexpression tobacco lines

The plasmid containing CaMV35Spro::*AtGASA6* and CaMV35Spro::*nptII* was transformed into the *Agrobacterium tumefaciens* strain LBA4404. 0.5×0.5 cm leaf explants were isolated from 6-week old *Nicotiana tabacum* aseptic seedlings and were co-cultivated with positive *Agrobacterium tumefaciens* colonies according to Horsch et al.1985 (Horsch et al. 1985). After co-cultivation under dark for three days, the treated explants were transferred to shooting medium with selection [1/2 MS + 2 mg/L 6-Benzylaminopurine (6-BA) + 0.1 mg/L (naphthalene-1-acetic acid) NAA + 50 mg/L kanamycin (Kan)+ 3% (W/V) sucrose + 0.8% agar]. After six weeks, well-developed shoots were transferred to the rooting medium (1/2 MS + 0.2 mg/L NAA + 30 mg/L Kan + 1.5% sucrose + 0.8% agar). Observations were conducted at different selective stages from explant inoculation to rooting. Putative transgenic T_0_ plants were transferred to soil and grew in a greenhouse.

### Regeneration of *AtGASA6* overexpression tobacco leaf explants

Seeds of *AtGASA6* overexpression tobacco lines were surface-sterilized in 1% sodium hypochlorite solution for 20 min. Then the seeds were rinsed in sterile water 4-5 times and germinated *in vitro* on the MS medium containing 3% sucrose and solidified with 0.8% agar (Murashige and Skoog 1962). Plants grew in a controlled-environmental growth chamber at 24°C and about 70% humidity with a 16-h-light/8-h-dark cycle. Six weeks after germination, 1 cm^2^ leaf discs were isolated from fully expanded leaves and used as explants. Explants were put on shooting media (1/2 MS + 2 mg/L 6-BA + 0.1 mg/L NAA + 0.8% agar) containing different concentration of sucrose (3%, 6%, 9%, and 0%). After six weeks, well-developed shoots were transferred to the rooting medium without sucrose (1/2 MS + 0.2 mg/L NAA + 0.8% agar). Observations were conducted at different selective stages from explant inoculation to rooting.

### Growth response of *AtGASA6* overexpression tobacco seedlings on sugar-free MS medium

The seeds of *AtGASA6* overexpression tobacco lines were sterilized according to the above method. Sterilized seeds were put on MS medium without sucrose. Shooting and rooting of seeds were observed.

### Plant transformation and regeneration to create *AtGAI* overexpression tobacco lines

The plasmid containing CaMV35Spro::*AtGAI* and CaMV35Spro::*AtGASA6* was transformed into tobacco according to the method of *AtGASA6* transformation. After co-cultivation under dark for three days, the treated explants were transferred to sugar-free shooting medium [1/2 MS + 2 mg/L 6-BA + 0.1 mg/L NAA + 0.8% agar]. After six weeks, well-developed shoots were transferred to the sugar-free rooting medium (1/2 MS + 0.2 mg/L NAA + 30 mg/L Kan + 0.8% agar). Observations were conducted at different selective stages from explant inoculation to rooting. Putative transgenic T_0_ plants were transferred to soil and grew in a greenhouse.

### RT-PCR and qRT-PCR

Total RNA was extracted from leaves using Trizol reagent according to the users’ manual. cDNA was synthesized using the first-strand cDNA synthesis kit (Tiangen, Beijing, China). Forward primer 5’-CGCTCTAGAGATGGCCAAACTCATAACTTC-3’ and reverse primer 5’-ATAGGATCCGATGGACATTTTGGTCCACC-3’ were used for *AtGASA6* amplification. Froward primer 5’-ACGACCAGCAAGATCCAACCGAAG-3’ and reverse primer 5’-ACGGAACAGGAATGGTTAAGGCTGG-3’ were used for *Actin* amplification. For qRT-PCR, samples were performed in triplicate using SYBR Green Master Mix on a Light Cycler 480 (Roche, Penzberg, Upper Bavaria, Germany). Relative levels of transcript were normalized to the level of *Actin* in each sample. Data shown are relative transcript abundance compared with that of control.

### Statistical analysis

For all statistical analyses, data are shown as means and standard errors (Mean±SE). Statistical significance was determined using the two-tailed Student’s *t*-test. For all tests, a p-value of less than 0.05 was considered statistically significant. p < 0.05, p < 0.01, p < 0.001, N.S., not significant versus control groups.

## Results

### Generation of *AtGASA6* overexpression tobacco lines

In our previous work, overexpression of *AtGASA6* was found to help transgenic *Arabidopsis* seeds survive on a medium with a high concentration of sugar (Zhong et al. 2015). To verify its function on sucrose metabolism, the overexpression cassette of *AtGASA6* (*p35S::AtGASA6*) (Figure 1a) was delivered into leaf explants of *Nicotiana tabacum* using the *Agrobacterium*-mediated transformation method. In the construct, *NPTII* (*p35S::NPTII*) was used as a selectable marker. The uninfected leaf explants were used as the negative control. After transformation, the infected leaf explants were moved to the shooting medium which contained 50 mg/L Kan. The infected leaf explants were subcultured every two weeks until shoots were generated. During the shooting process, putative transgenic explants grew green and viable shoots (Figure 1b) while uninfected leaf explants turned pale, necrotic, and eventually died (Figure 1c). Generated shoots were further transferred to the rooting medium containing 30 mg/L Kan and also subcultured every two weeks (Figure 1d). Putative transgenic shoots could grow roots while negative ones could not (Figure 1d).

**Figure 1.**
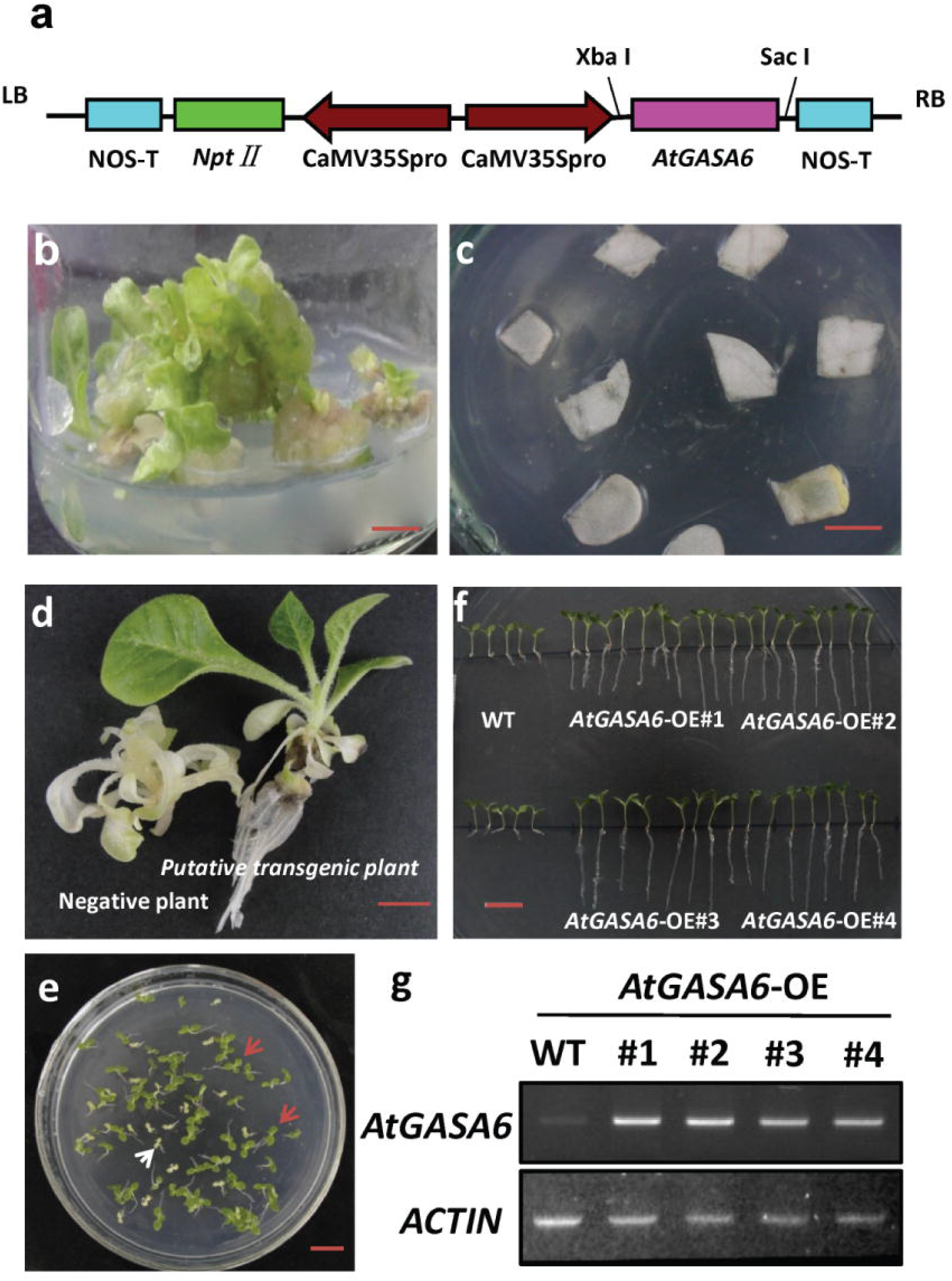
Generation of *AtGASA6* overexpression tobacco lines. (a) Schematic diagram of the binary vector used for transformation. The binary vector pBIN19 is used as the backbone. Both *AtGASA6* and *NPTII* genes are driven by the constitutive *CaMV 35S* promoter (CaMV35Spro). The two expression cassettes are in reverse orientations. NOS-T: nopaline synthase promoter/terminator; LB: left border; RB: right border. (b) Putatively transformed leaf explants are growing shoots on shooting medium. Bar=1 cm. (c) Untransformed leaf explants are turning pale on shooting medium. Bar=1 cm. (d) The putative transgenic shoot (right) grows roots while the negative shoot (left) cannot grow roots and is turning pale on rooting medium. Bar=1 cm. (e) The ratio of normal and pale plants is 3:1 when T_1_ generation seedlings of *AtGASA6* transgenic lines are grown on MS medium containing 50 mg/L Kan. Bar=1 cm. (f) Young seedlings of T_2_ generation of the four *AtGASA6* transgenic lines grow normally on MS medium containing 50 mg/L Kan while the growth of WT seedlings is suppressed. Bar=1 cm. (g) The transcription of *AtGASA6* is detected in leaves of each of the four overexpression lines (T_2_ generation) in RT-PCR analysis. *Actin* is used as a house-keeping gene.

Four plants, *AtGASA6*-OE#1, *AtGASA6*-OE#2, *AtGASA6*-OE#3, and *AtGASA6*-OE#4, with both shoots and roots, were created and considered as putative transgenic lines. Seeds (T_1_ generation) of each line were harvested and grown on the MS medium containing 50 mg/L Kan (Figure 1e). The ratio of normal and pale plants was 3:1 which was consistent with Mendel’s Law of Segregation (Figure 1e). Seeds (T_2_ generation) of the T_1_ generation were further grown on MS medium containing 50 mg/L Kan (Figure 1f). Young seedlings of the four *AtGASA6* transgenic lines grew normally on MS medium containing Kan while the growth of WT seedlings was suppressed (Figure 1f). The T_1_ generation plants whose seeds had no segregation on the MS medium containing Kan were considered as homozygotes and their seeds (T_2_ generation) were used for all the following experiments in this work. In the RT-PCR analysis, a bright band of *AtGASA6* was found in leaves of each of the four transgenic lines (Figure 1g), which indicates the successful overexpression of *AtGASA6*.

### Leaf explants of *AtGASA6* overexpression tobacco lines regenerate on sugar-free media

To investigate whether *AtGASA6* could act as a selectable marker that enables transformed plants to survive on media with or without sucrose, leaf explants of *AtGASA6* overexpression lines were put on the shooting media with 3% (normal), 6%, 9%, and 0% sucrose. The leaf explants of WT were used as the control. After four weeks on the shooting medium with 3% and 6% sucrose, the leaf explants of both *AtGASA6* overexpression lines and WT generated shoots (Figure 2a-b). On the shooting medium with 9% sucrose, the leaf explants of both *AtGASA6* overexpression lines and WT generated brown shoots that were unviable (Figure 2c). These data indicate that 3% and 6% sucrose cannot distinguish *AtGASA6* transformed plants from WT, and the higher (9%) sucrose concentration hampers shoot regeneration.

**Figure 2.**
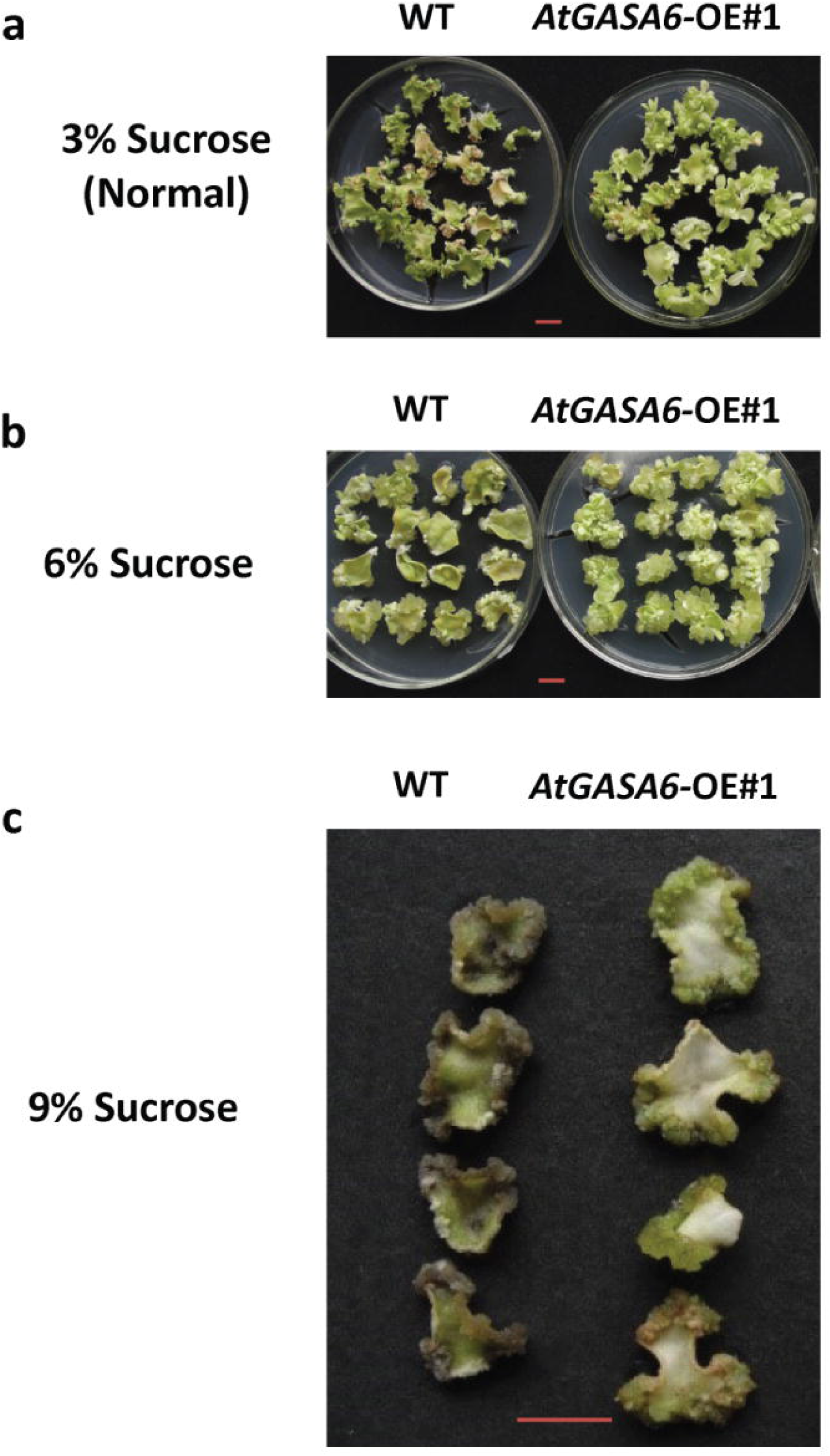
Testing of sucrose concentration on which explants of *AtGASA6* overexpression lines can regenerate while explants of WT cannot. (a-b) Calli are induced and shoots are subsequently regenerated from leaf explants of both *AtGASA6* overexpression lines and WT on shooting media with 3% (a) and 6% (b) sucrose. Bar=1 cm. (c) Leaf explants of both *AtGASA6* overexpression lines and WT is browning and no shoot regeneration is observed on shooting medium with 9% sucrose. Bar=1 cm. Considering the similar phenotype, only the picture of overexpression line *AtGASA6*-OE #1 is shown.

Attractively, on the shooting medium without sucrose, approximately three shoots were observed in each leaf explant of *AtGASA6* overexpression lines while no shoots were found in any WT leaf explants (Figure 3a-b). The shoots of *AtGASA6* overexpression lines were further transferred to the rooting medium without sucrose and each shoot regenerated roots (Figure 3c). Collectively, leaf explants of *AtGASA6* overexpression lines could regenerate shoots on the sugar-free shooting medium while leaf explants of WT had no shoot regeneration. This indicates that *AtGASA6* could be used as a selectable marker on the sugar-free shooting medium.

**Figure 3.**
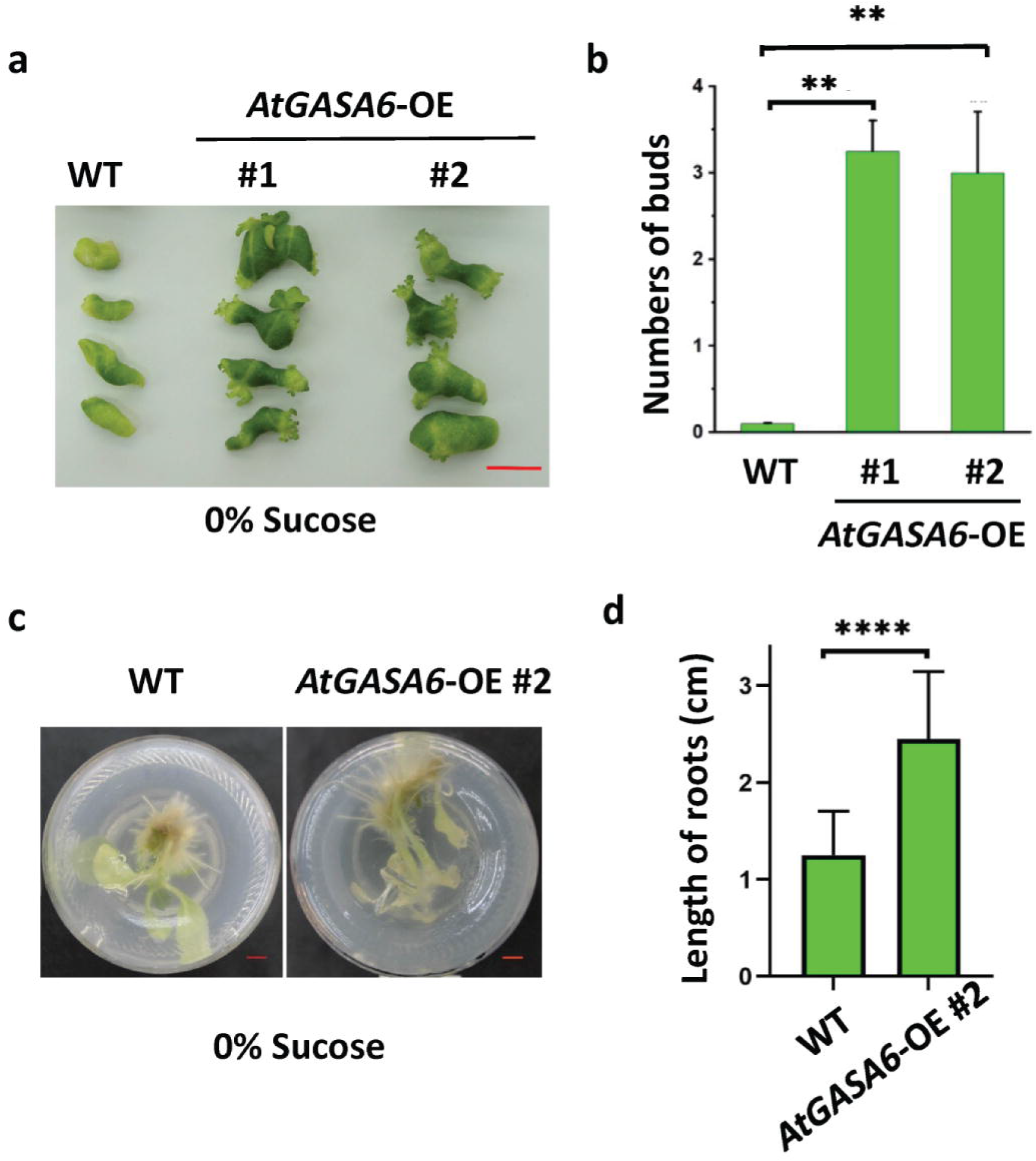
Leaf explants of *AtGASA6* overexpression lines regenerate shoots and roots on sugar-free shooting and rooting media, respectively, while leaf explants of WT cannot. (a) Leaf explants of *AtGASA6* overexpression lines regenerate shoots on sugar-free shooting medium while leaf explants of WT cannot. Bar=1cm. (b) Almost each leaf explant of *AtGASA6* overexpression lines has three regenerated shoots while each leaf explant has zero shoot on sugar-free rooting medium. (c) Shoots of *AtGASA6* overexpression lines can grow roots normally on sugar-free rooting medium. Bar=1cm. Considering the similar phenotype, only the pictures of overexpression lines AtGASA6-OE #1 and/or #2 are shown.

### *AtGASA6* enables seed selection

In addition to the selection on callus stage, selection on seed is another key selection method in plant transformation. To investigate whether *AtGASA6* could be used as a selectable marker in plant species that use seeds for selection (e.g. *Arabidopsis*), the seeds of *AtGASA6* overexpression lines were germinated on the MS medium without sucrose. The seeds of WT were used as the negative control. One week later, green seedlings were germinated from seeds of all *AtGASA6* overexpression lines. However, only yellow seedlings were grown from WT seeds (Figure 4a). Consistently, the relative chlorophyll content in all *AtGASA6* overexpression lines was almost 1.5 folds higher than that in WT seedlings (Figure 4b). A large number of long roots were observed from each seedling of all the *AtGASA6* overexpression lines while only a few short roots were found from WT seedlings (Figure 4c). Moreover, the length and the number of leaves in *AtGASA6* overexpression line seedlings were much larger than those of WT seedlings (Figure 4d). Due to the bad health condition, the WT seedlings died eventually. The fact that *AtGASA6* overexpression seeds survive on the sugar-free MS medium indicates that *AtGASA6* could be used as a selectable marker on seed selection.

**Figure 4.**
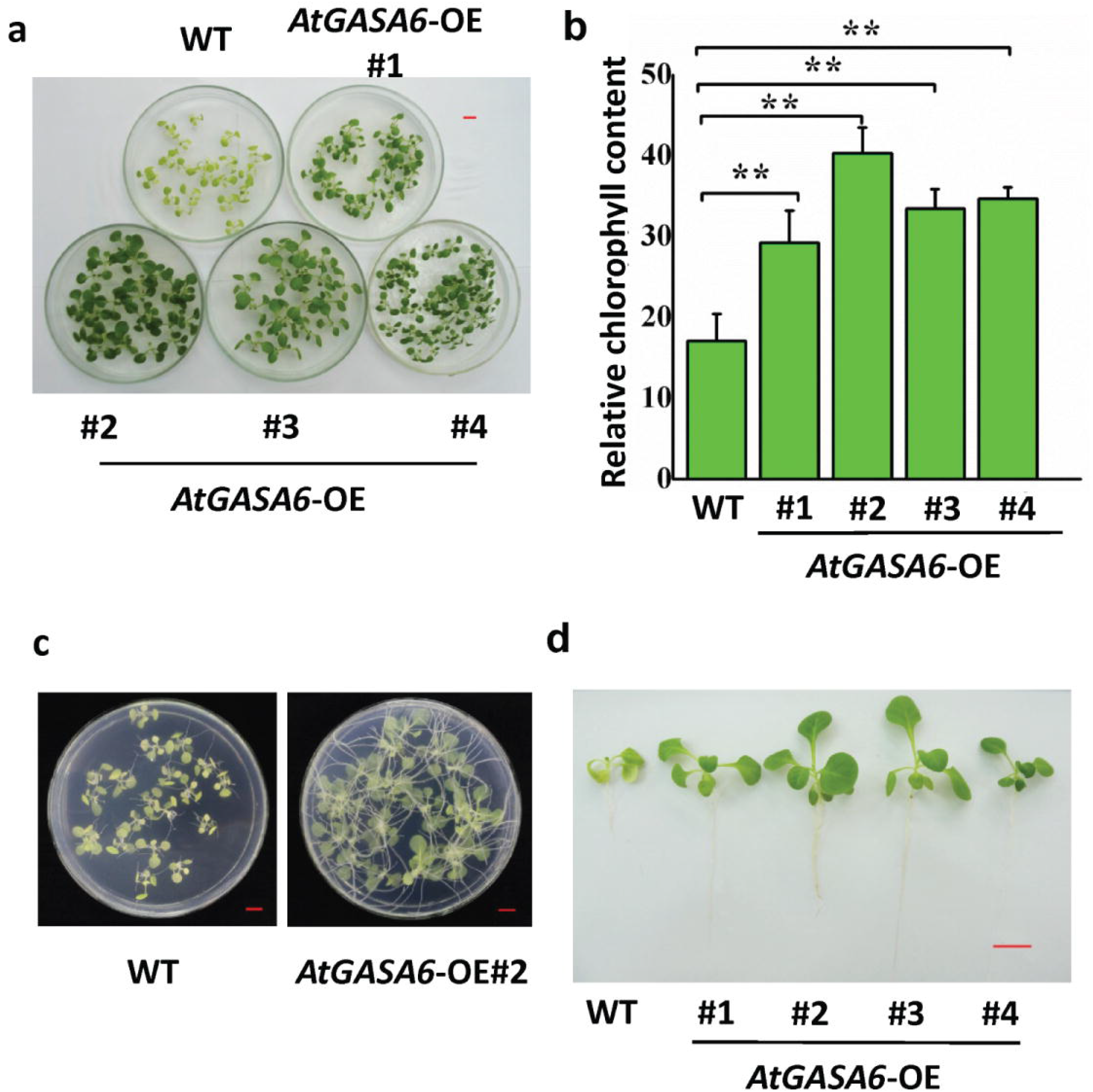
Growth response of *AtGASA6* overexpression tobacco seedlings on sugar-free MS medium. (a) Seedlings of *AtGASA6* overexpression lines have normal shoots while seedlings of WT have pale shoots on the sugar-free MS medium. Bar=1 cm. (b) Seedlings of *AtGASA6* overexpression lines have normal roots while seedlings of WT have less and short roots on the sugar-free MS medium. Considering the similar phenotype, only the picture of overexpression line *AtGASA6*-OE #2 is shown. Bar=1 cm. (c) Seedlings of *AtGASA6* overexpression lines have normal shoots and roots while seedlings of WT have abnormal shoots and roots on the sugar-free MS medium. Bar=1 cm. (d) The chlorophyll content of *AtGASA6* overexpression lines is significantly higher than that of WT on the sugar-free MS medium.

### No developmental defects are observed in *AtGASA6* overexpression plants

In addition to the assistance in selection, no influence on the development of transgenic plants is a key criterion of a successful selectable marker gene. The morphologic performance between *AtGASA6* overexpression lines and WT was compared. During the seedling stage, the leaf, stem, and root size had no obvious variations between *AtGASA6* overexpression lines and WT on MS medium with 3% (the normal concentration) sucrose (Figure 5a). Similarly, there was also no apparent difference in the leaf and stem size between *AtGASA6* overexpression lines and WT in the soil during the vegetative phase (Figure 5b). In addition to the leaves and stems, the flowers had no obvious variations between *AtGASA6* overexpression lines and WT during the reproductive growth stage (Figure 5c-d). Moreover, the size and number of capsules also had no apparent difference between *AtGASA6* overexpression lines and WT (Figure 5e-f). Taken together, the overexpression of *AtGASA6* has no observable influence on the development of transgenic plants.

**Figure 5.**
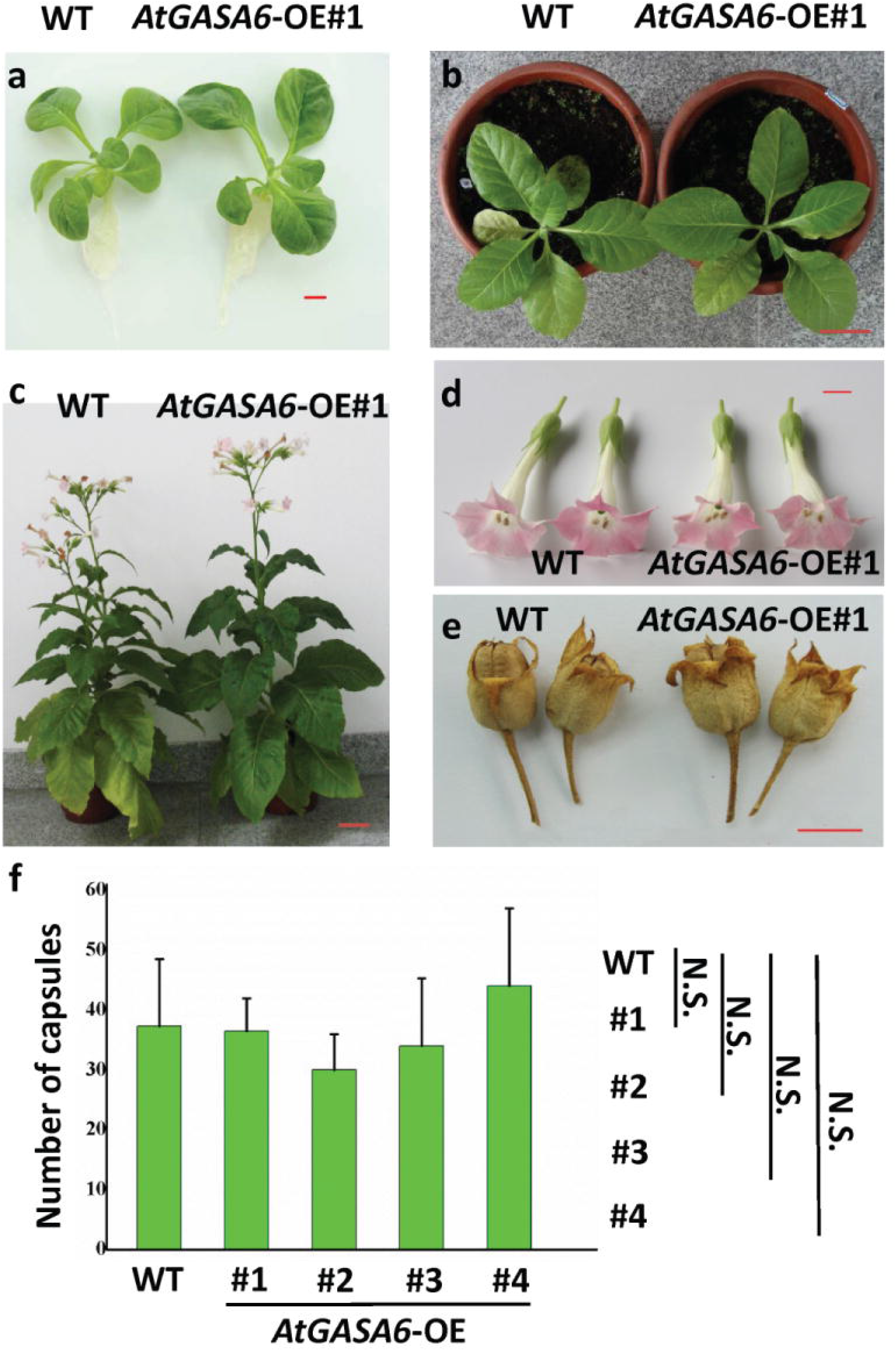
No growth defects are observed in *AtGASA6* overexpression lines. (a-c) Morphologic comparison between *AtGASA6* overexpression lines and WT growing on MS medium in a growth chamber (a) and in soil in a greenhouse (b-c). Bar=1 cm. (d-e) Morphologic comparison of flowers (d) and capsules (e) between *AtGASA6* overexpression lines and WT. Bar=1 cm. Considering the similar phenotype, only the pictures of overexpression line *AtGASA6*-OE #1 are shown. (f) There is no significant difference in the number of capsules between *AtGASA6* overexpression lines and WT. N.S., no significant difference.

### *AtGASA6* assisted gene transformation in tobacco

To verify the feasibility as a selectable marker in gene transformation, *AtGASA6* was co-delivered with *AtGAI*, which controls plant size, into *Nicotiana tabacum* using the *Agrobacterium*-mediated transformation method. In this transformation, the overexpression cassettes of *AtGASA6* (CaMV35Spro::*AtGASA6*:NOS-T) and *AtGAI* (CaMV35Spro::*AtGAI*: NOS-T) were comprised in a single construct in reverse orientations. Construct merely containing the *AtGASA6* overexpression cassette (CaMV35Spro::*AtGASA6*:NOS-T) was used as a negative control (Figure 6a). After transformation, explants were transferred to the sugar-free shooting medium, and regenerated shoots were subsequently transferred to the sugar-free rooting medium. Three putative *AtGAI* lines (AtGAI-OE#1, AtGAI-OE#2, and AtGAI-OE#3) and one negative control line were obtained (Figure 6b). In the qRT-PCR analyses, *AtGASA6* had relative transcription levels of 1.0-1.35 in the leaves of all putative *AtGAI* and negative control lines (Figure 6c). Moreover, *AtGAI* had relative transcription levels of 5-10 in leaves of all putative *AtGAI* lines and had no transcription in the negative control lines (Figure 6d). These data demonstrate that *AtGASA6* and *AtGAI* have been successfully transformed and expressed. Consistent with the overexpression of *AtGAI*, the lengths of the third leaves of all the three *AtGAI* overexpression lines were 1.9 ∼2.8 cm, which was significantly shorter than the 4.3 ± 0.3 cm of that of the negative control. Similarly, the lengths of the fourth, fifth, and sixth leaves, were two, three, and four times shorter than those of the negative control, respectively (Figure 6b, Table 1). These data collectively demonstrated that transgenic tobacco lines of *AtGAI* had been successfully generated using *AtGASA6* as a selectable marker gene.

**Figure 6.**
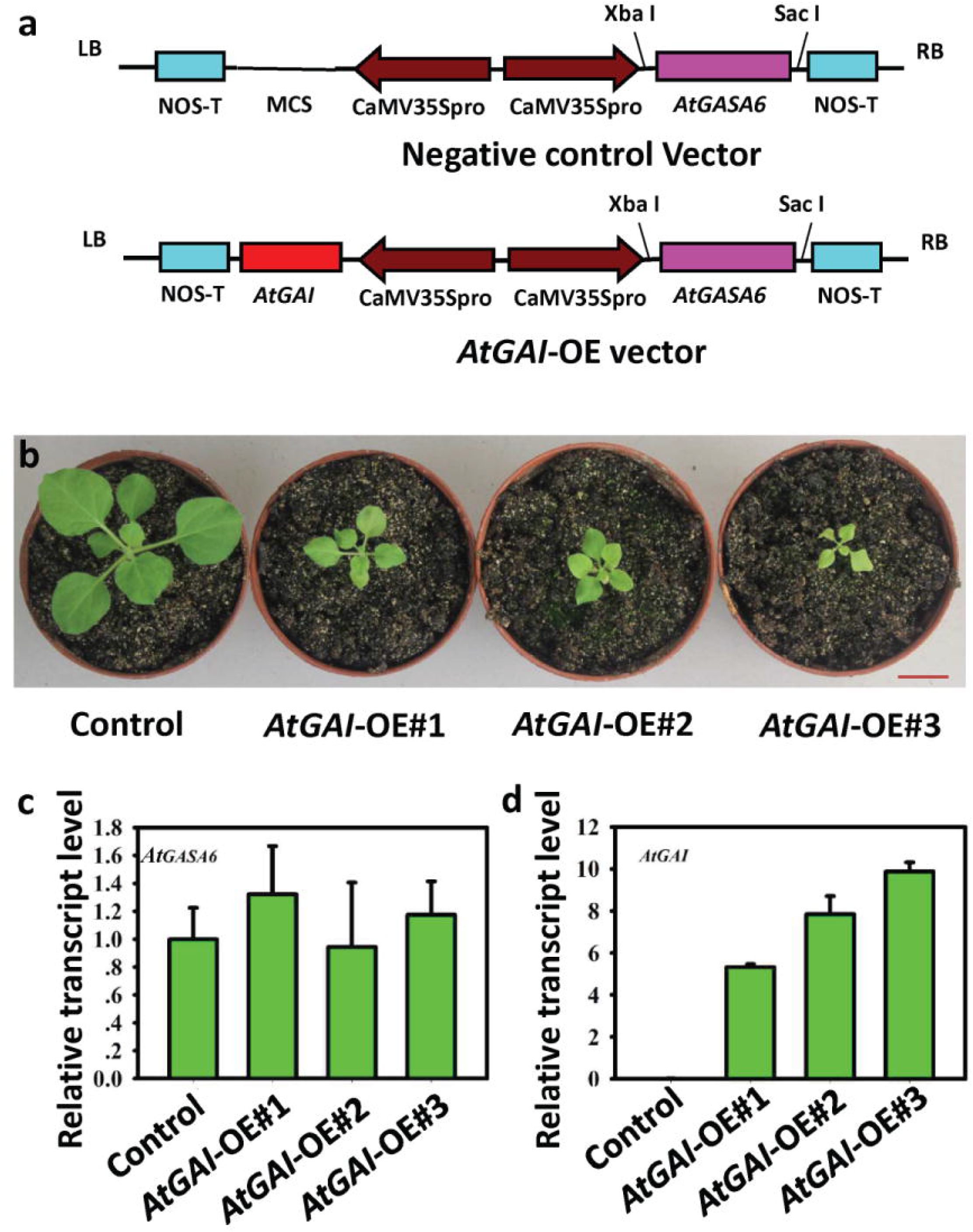
Generation of tobacco transgenic lines using *AtGASA6* as a selectable marker. (a) Schematic diagrams of the binary vectors used for transformation. The binary vector pBIN19 is used as the backbone of the vector. Both *AtGASA6* and *AtGAI* genes are driven by the constitutive *CaMV 35S promoter* (CaMV35Spro). The two expression cassettes are in reverse orientations. NOS-T: nopaline synthase promoter/terminator; LB: left border; RB: right border; MCS: multiple cloning site. The negative control vector contains the *AtGASA6* expression cassette but not the *AtGAI* expression cassette. OE, overexpression. (b) The plant size of *AtGAI* overexpression lines generated using *AtGASA6* as a selectable marker on the sugar-free shooting and rooting media is smaller than that of the negative control line. AtGAI-OE#1, AtGAI-OE#2 and AtGAI-OE#3 are three overexpression lines of *AtGAI* gene. Control is one of positive lines of the negative control vector. Bar= 1 cm. (c) The overexpression of *AtGASA6* in the leaves of both *AtGAI* overexpression lines and the negative control line is detected in qRT-PCR analysis. (d) The overexpression of *AtGAI* is detected in leaves of *AtGAI* overexpression lines while not in leaves of the negative control line in qRT-PCR analysis.

**Table 1.** The leaf length of *AtGAI* overexpression tobacco lines. The leaf length covers both petiole and leaf. The data are the means ± SEs. n=3 and n=15 for the control and *AtGAI* overexpression lines, respectively. Asterisks indicate statistically significant differences at P < 0.05 (*) and P< 0.01 (**) between the overexpression lines and WT by one-way ANOVA in the SPSS program.

| Plant genotype | Leaf length (cm) |  |  |  |
| --- | --- | --- | --- | --- |
|  | The third leaf | The fourth leaf | The fifth leaf | The sixth leaf |
| **Control** | 4.3 +/- 0.3 | 5.4 +/- 0.2 | 6.8 +/- 0.4 | 6.6 +/- 0.5 |
| *AtGAI-OE#1* | 2.8 +/- 0.2** | 2.5 +/- 0.3** | 2.1 +/- 0.2** | 1.5 +/- 0.1** |
| *AtGAI-OE#2* | 2.4 +/- 0.1** | 2.2 +/- 0.2** | 2.0 +/- 0.3** | 1.8 +/- 0.4** |
| *AtGAI-OE#3* | 1.9 +/- 0.3** | 1.7 +/- 0.4** | 1.4 +/- 0.2** | 1.2 +/- 0.3** |

## Discussion

### Advantages of using *AtGASA6* as a selectable marker

As a selectable marker, *AtGASA6* has four advantages. First, *AtGASA6* is a plant-derived gene and will not trigger human health issues. In contrast, most of the traditional selectable marker genes, such as *nptII* and *bar*, are from bacteria and hence raise severe human health concerns. Second, the *AtGASA6* assisted selection system only needs sugar-free shooting medium during selection and does not need any antibiotics that always are expensive and toxic, which makes this system low-cost and safe. Third, transgenic plants could be distinguished unambiguously from non-transgenic plants both at the shooting (Figure 3) and young seedling stages (Figure 4). This indicates that *AtGASA6* could be used as a selectable marker in both plant cell/tissue and floral dipping, which are two major plant transformation approaches. Fourth, *AtGASA6* transgenic plants have no obvious developmental defects. No morphological abnormalities is the prerequisite of a successful selectable marker. The transformants produced using Kan selection always display retarded growth even after being transplanted to soil. This phenomenon is attributed to the inhibitory effect of Kan on transgenic plants (Lindsey and Gallois 1990). In this study, *AtGASA6* overexpression tobacco lines were morphologically normal (Figure 5). All the above four advantages enable *AtGASA6* a promising selectable marker in plant transformation.

### *GASA6* is widely distributed in plants

The application of *GASA6* would be extensively broadened if every plant species could use the *GASA6* gene of its own. To address this question, the distribution of *GASA6* in plants was analyzed. In the BLASTP search using the protein sequence of *AtGASA6* as a query, homologs were found in a broad range of plant species. These species include major cash crops, such as *Raphanus sativus*, *Camelina sativa*, *Brassica napus*, *Manihot esculenta*, and *Punica granatum*. All the homologs had 100% of query cover and had protein identity percentages from 68.27% to 90.10% with *AtGASA6* (Figure S1). The EggNOG database and SMART were further used to analyze the functional domains among *AtGASA6* homologs. In the EggNOG analysis, the GASA functional domain (from 42Q to 101P), which might exert gibberellin-regulated function, was shared by all homologs (Figure S2a). In the SMART analysis, conserved signal peptide and transmembrane domain were shared among all homologs (Figure S2b). The conserved functional domains indicate that *GASA6* homologs in most plant species might exert a similar molecular function with *AtGASA6*. The wide distribution of homologs and the similar function opens the possibility that most plant species could use their native *GASA6* gene as a selectable marker in transformation.

### Improvement of the *AtGASA6*-assisted selection system

The constitutive *CaMV* 35S promoter was used to drive the overexpression of *AtGASA6* in this work. Although no obvious morphological variations were observed in transgenic tobacco plants compared with WT, the constitutive overexpression of *AtGASA6* might increase the risk of developmental defects in other plant species. Callus-specific or seed-specific expression promoters, such as the promotor of cysteine proteinase gene (*OsCSP*) in rice (Ray et al. 2004) and *P1830* in maize (Yu et al. 2019), could be used to drive the expression of *AtGASA6*. The callus or seed specifical expression of *AtGASA6* might reduce the risk of developmental defects. In recent years, emerging techniques have been developed to remove selectable markers in transformants. For example, the FLP-CRE system could remove the selectable marker gene by a further cross to induce Cre expression (Huang et al. 2020; Zhao et al. 2019). Using the FLP-CRE system, *AtGASA6* could also be removed from transgenic progenies. In conclusion, *AtGASA6* could be a promising plant-derived, non-antibiotic and widely distributed selectable marker in plant transformation.

## Author contributions

S.Z., X.W., and Y.L. designed experiments. Y.L., J.G. performed experiments. Y.L., J.G., S.Z., and X.W. performed data analysis. Y.L. wrote the manuscript. All authors approved the submission.

## Acknowledgments

This work is supported by the National Natural Science Foundation of China (31870301, 31370350 for S.Z). We thank Dr. Guangbin Luo (University of Copenhagen, Denmark) for polishing the manuscript.

## Supporting information

**Figure S1.**
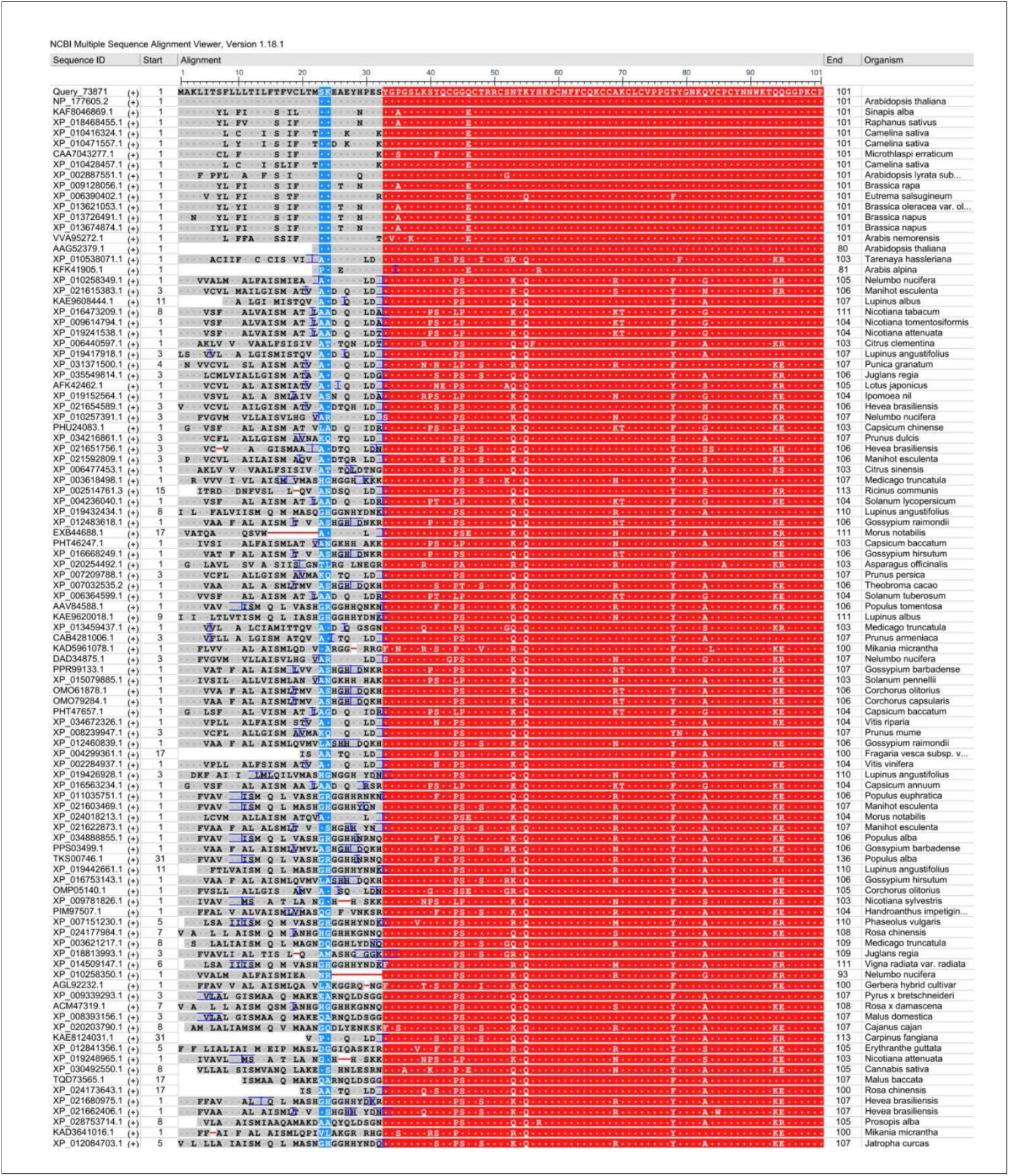
Homologous of *AtGASA6* are found in a variety of plant species in NCBI BLAST.

**Figure S2.**
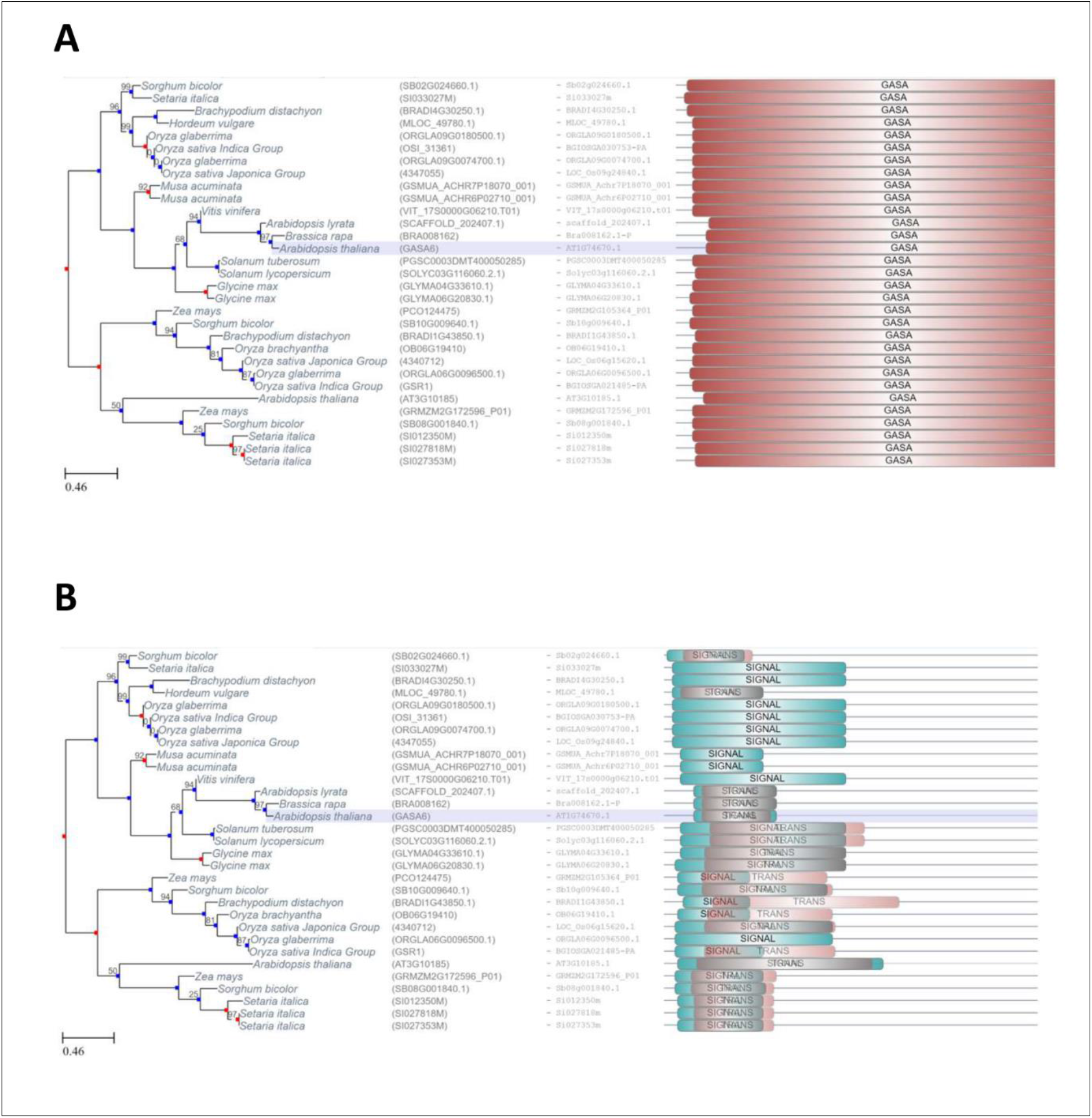
Homologous of *GASA6* among different plant species have conserved functional domains. (A) Homologous of GASA6 share the conserved GASA6 domain in the PFAM domain analysis. (B) Homologous of GASA6 share the conserved signal and trans domains in the functional domain analysis.

