## Supplementary figures and images for "A novel non-antibiotic selectable marker *GASA6* for plant transformation"

### FigureS1.tif

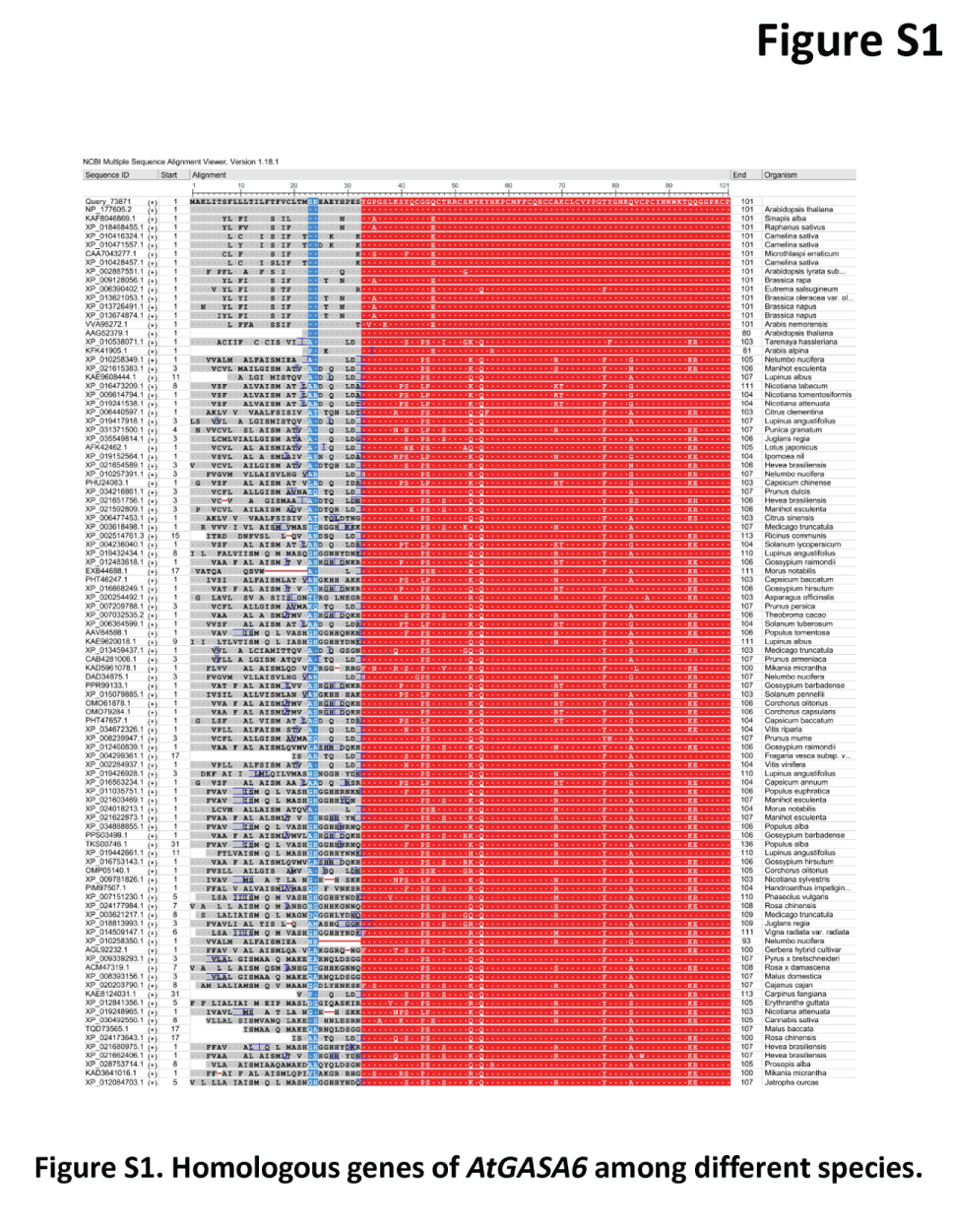

### FigureS2.tif

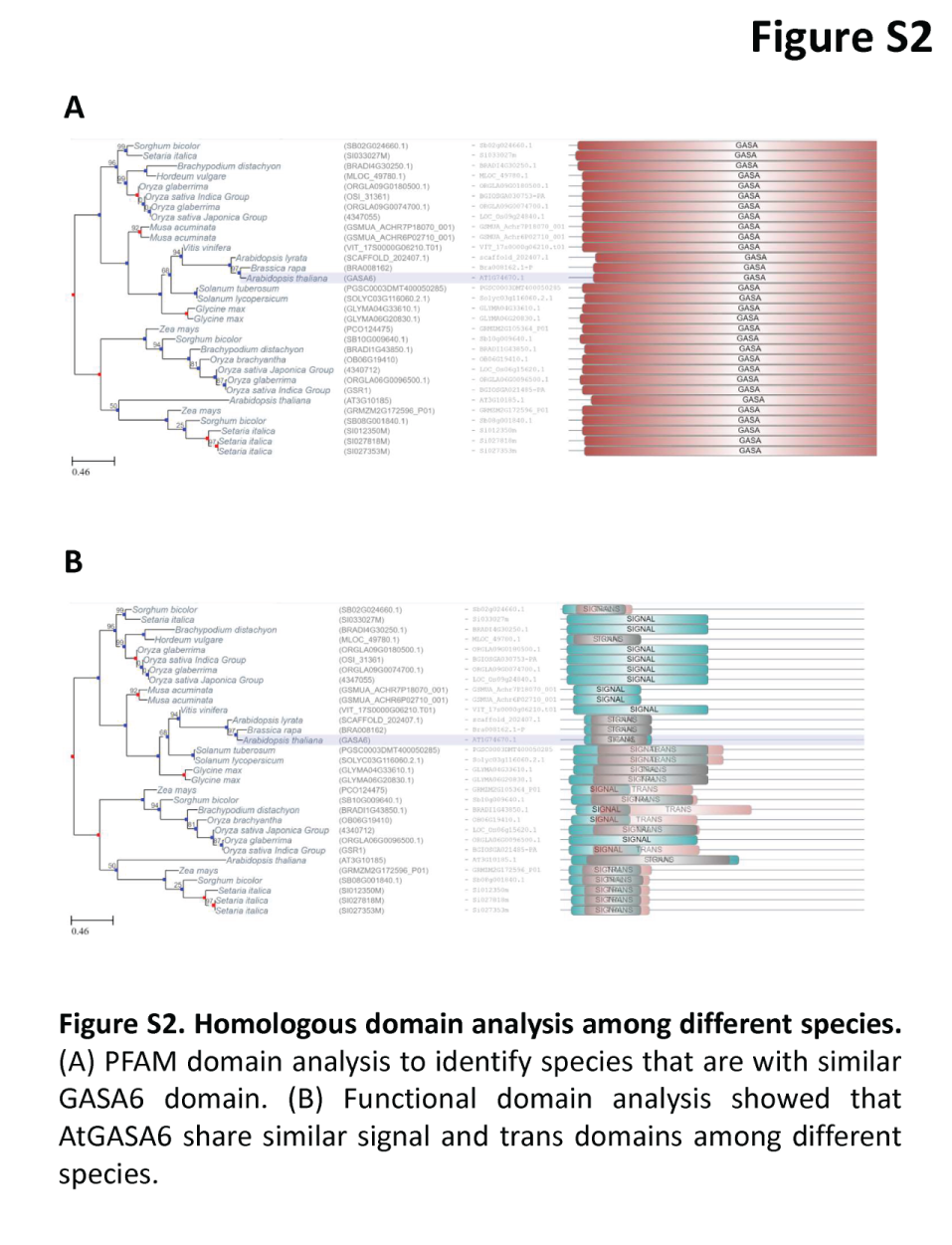
